# Word stress representations are language-specific: evidence from event-related brain potentials

**DOI:** 10.1101/442129

**Authors:** Ferenc Honbolygó, Andrea Kóbor, Borbála German, Valéria Csépe

## Abstract

Understanding speech at the basic levels entails the simultaneous and independent processing of phonemic and prosodic features. While it is well-established that phoneme perception relies on language-specific long-term traces, it is unclear if the processing of prosodic features similarly involves language-specific representations. In the present study, we investigated the processing of a specific prosodic feature, word stress, using the method of event-related brain potentials (ERPs) employing a cross-linguistic approach. Hungarian participants heard disyllabic pseudowords stressed either on the first (legal stress) or on the second (illegal stress) syllable, pronounced either by a Hungarian or a German speaker. Results obtained using a data-driven ERP analysis methodology showed that all pseudowords in the deviant position elicited an Early Differentiating Negativity (EDN) and a Mismatch Negativity (MMN) component, except for the Hungarian pseudowords stressed on the first syllable. This suggests that Hungarian listeners did not process the native legal stress pattern as deviant, but the same stress pattern with a non-native accent was processed as deviant. This implies that the processing of word stress was based on language-specific long-term memory traces.

## Introduction

Understanding speech requires the simultaneous processing of segmental and suprasegmental (prosodic) information, suggested to be based on parallel and independent processing mechanisms (Poeppel, 2014), and possibly relying on separate neural pathways (Bornkessel-Schlesewsky & Schlesewsky, 2013). Among the available prosodic information, word stress is a relative emphasis given to certain syllables within words or to certain words in sentences (for review, see Kager, 2007). Word stress can emphasize or separate certain parts of the speech stream; thus, it potentially contributes to the segmentation of continuous speech into words (Cutler & Norris, 1988). However, languages differ in several aspects regarding word stress: in the position of the stressed syllable within multisyllabic words (initial, final, penultimate, etc.); in the variability of the stressed syllable’s position (free or fixed); and whether stress can distinguish lexical meaning (contrastive or non-contrastive). Consequently, the language processing system of individuals has to adapt to the specific features of word stress of a given language. In the present study, we investigated whether stress related representations can be considered as language-specific, and whether we find any evidence of language-specific processing at the neurocognitive level.

Previous studies investigating the neurocognitive basis of word stress processing studied mostly the Mismatch Negativity (MMN) event-related brain potential (ERP) component. The MMN is an auditory ERP component with a negative polarity and a frontocentral voltage maximum, and has been an exceptionally useful tool in studying linguistic processing (for review, see Näätänen, Paavilainen, Rinne, & Alho, 2007). The MMN is usually elicited in passive oddball paradigms where frequently repeated standard stimuli are interspersed by rarely repeated deviant stimuli differing from the standard in some discriminable features.

The MMN appears 100-250 ms after the onset of the change and can be elicited in the absence of participants’ attention. The MMN is currently interpreted as a brain electrical correlate of the mainly pre-attentive detection of violation of simple or complex regularities (Winkler, Denham, & Nelken, 2009).

A more complex model for the MMN’s emergence is proposed in the AERS (Auditory Event Representation System) predictive coding model suggested by Winkler and Schröger (2015). According to the model, acoustic information reaching the auditory system undergoes an initial sound analysis which extract the proto-features of sounds (for speech, these are speech-related acoustic-phonetic information, such as voicing, formant transitions, fundamental frequency, intensity, etc.). This is followed by binding the separate features to unitary sensory memory representations (for speech, these are phonetic representations like phonemes, prosodic features, etc.). These representations are then compared against the predictions derived from the model of the acoustical environment. The predictive model stores representations of the current regularities of the acoustical environment and it is also affected by long-term experience, which is available for native linguistic representations (speech sounds, prosodic patterns, words, etc.). The predictive model is the basis of the mechanisms comparing incoming information with the regularity representations. If predictions from the model fail, the model needs to be corrected via an updating process, which is reflected by the MMN.

There are a number of studies that applied the MMN paradigm to study the processing of word stress. Weber et al. (2004) found that German adults showed an MMN to the word ‘baba’ (‘baby’ in English) in the deviant position both in the trochaic form (stress on the first syllable, i.e., /ba⍰ba/) and in the iambic form (stress on the second syllable, i.e., /baba⍰/), indicating that both stress patterns were equally well discriminated.

Ylinen et al. (2009) studied the effect of prosodic familiarity in the case of Finnish words and pseudowords with familiar (trochaic, i.e., stress on the first syllable) and unfamiliar (iambic, i.e., stress on the second syllable) stress patterns. They used the pseudowords with the familiar stress pattern as standard and investigated the MMNs elicited by pseudowords with unfamiliar stress pattern, words with familiar stress pattern, and words with unfamiliar stress pattern. According to the results, the pseudowords with unfamiliar stress pattern elicited two MMNs related to the first and second syllables of utterances and were interpreted as the detection of the unexpected lack of stress on the first syllable and the detection of unexpected presence of stress on the second syllable, respectively. Words with familiar stress pattern elicited a single MMN in mid-latency time window, which was interpreted as the detection of the change of the lexical status of the word, and words with unfamiliar stress pattern elicited a single MMN related to the presence of stress on the second syllable. These results demonstrated that the familiarity of the words modulated the processing of stress pattern, and also that the familiarity of stress pattern modulated the processing of words.

Zora, Heldner, & Schwarz (2016) investigated stress processing in Turkish. In the study, the authors presented word and pseudoword stimuli differing in their stress pattern, with stimuli with iambic stress pattern being always standards and stimuli with trochaic pattern being deviants. They manipulated the acoustic features contributing to stress (f0, spectral emphasis, duration, all features combined), in order to investigate to what extent they are utilized for lexical access. Results showed that changes in prosodic features elicited ERPs both in words and pseudowords but there were processing differences between the prosodic features. The most prominent feature was the f0 eliciting an MMN in words but a frontal P3a in pseudowords, and consequently, was claimed to be lexically specified in Turkish by the authors. Results also indicated that the individual prosodic features rather than combined features differentiated language-related effects, since the combined features elicited similar MMN in both words and pseudowords.

The same authors (Zora, Schwarz, & Heldner, 2015) investigated also the contribution of two specific features, f0 and intensity to stress processing in English. In the study, participants heard stress minimal pairs versions (i.e., the same word with trochaic and iambic stress pattern) of the word ‘upset’ and the pseudoword ‘ukfet’. Deviants – having trochaic stress compared to the standards having iambic stress – differed from the standards in f0, intensity, or the combination of the two. According to the results, all deviants elicited the MMN, although intensity change elicited a smaller MMNs in pseudowords than f0, confirming the importance of both f0 and intensity to determine words stress and contribute to lexical access.

Finally, in a study comparing the processing of duration-related stress in speech and music in English (Peter, Mcarthur, & Thompson, 2012), the authors found that in the case of speech, only the stress on the first syllable condition elicited an MMN, while in the case of music stimuli, both long-short and short-long patterns (the musical equivalent of stress on the first and stress on the second syllable) elicited an MMN, indicating a fundamental difference between speech and non-speech related processing.

The above studies established convincingly that the MMN can be reliably used to investigate the neurocognitive background of stress processing and demonstrated the sensitivity of this particular ERP component to acoustic, phonetic, and lexical effects. In a series of studies, we investigated word stress processing in Hungarian. Hungarian is an excellent target language for studying word stress representations because the stress pattern of disyllabic word is highly regular: stress always falls on the first syllable. Therefore, any changes in the stress pattern is considered as violating the regularity, leading to illegal stress patterns. First, it was demonstrated that a word with stress on the second syllable elicited two MMN components when contrasted with a word with stress on the first syllable (Honbolygó, Csépe, & Ragó, 2004). In a subsequent study (Honbolygó and Csépe, 2013), it was also shown that pseudowords with stress on the second syllable elicited two consecutive MMN components, while pseudowords with a familiar stress pattern in a deviant position did not elicit an MMN, suggesting that stress processing is modulated by top-down processes, supporting the importance of familiarity in word stress processing. This study was replicated using meaningful words (Garami, Ragó, Honbolygó, & Csépe, 2017), demonstrating that lexicality facilitates the processing of familiar stress patterns but not the unfamiliar ones. These studies provided further evidence that Hungarian speakers rely on long-term representations of the fixed Hungarian stress pattern that appear to contain a description of the expected word stress pattern characteristic for a given language at the phonological level. However, the language-specific nature of these representations has not been directly investigated in a cross-linguistic experiment so far; therefore, the present study specifically aimed to clarify this aspect.

The MMN paradigm has also been used to investigate language-specific processing of linguistic representations. It has been demonstrated in several studies (Dehaene-Lambertz, 1997; Kraus & Cheour, 2000; Näätänen et al., 1997; Winkler et al., 1999) that native language phonemes elicit enhanced MMN as compared to non-native phonemes, indicating the modulation of speech processing mechanisms by the exposure to native language. Chandrasekaran, Krishnan, & Gandour (2007) used the MMN paradigm to study the effect of language experience on the processing of pitch contour in Mandarin Chinese in a cross-linguistic study. According to the results, native Chinese speakers showed a larger MMN component to highly dissimilar tones than non-native (English) speakers, demonstrating that the MMN is modulated by the long-term experience with the native language.

In the present study, we also used the MMN paradigm to investigate the language-specific nature of word stress representations in native speakers of Hungarian. In the study, we used naturally produced, meaningless pseudowords having stress either on the first or the second syllable. Moreover, we created a native and a non-native (German) version of both types of pseudowords in order to study the ERP correlates of non-native stress processing. We assumed that listeners would detect preattentively if a pseudoword was pronounced in a native or non-native way, and only the native-like stimuli will activate the long-term stress representation. We expected that the activation of the long-term representation would be indicated by not recognizing the native legal stress patterns as deviant (as they do not differ from the expected stress pattern specified in the long-term representation). Based on these assumptions, we hypothesized that pseudowords stressed on the second syllable (illegal stress pattern in Hungarian) would elicit the MMN component irrespective of the language. However, we expected that pseudowords stressed on the first syllable (legal stress pattern in Hungarian) would elicit the MMN component only when the non-native version is presented, and no MMN would be elicited by the native legal pseudoword, because listeners would not detect this pattern as irregularity.

## Methods

### Participants

Thirty-two healthy young adults took part in the experiment. Two of them were excluded by reason of excessive artifacts (see EEG Recording and Pre-Processing). Therefore, thirty participants remained in the final sample (19 females). The participants, aged between 18 and 25 years (*M*_Age_ = 21.7 years, *SD* = 1.6 years), were native speakers of Hungarian who have not studied German as L2 according to a questionnaire screening their linguistic background (Kóbor et al., 2018). All of them were right-handed according to the Edinburgh Handedness Inventory revised version (Dragovic, 2004) (*M*_LQ_ = 90.4, *SD* = 12.9^1^). Most of the participants were undergraduate students enrolled in different universities in Budapest (*M*_Years of education_ = 15.1 years, *SD* = 1.6 years). They had normal or corrected-to-normal vision and normal hearing level according to the audiometry measurement. Prior to their inclusion in the study, participants provided informed consent to the procedure as approved by the Ethical Review Committee for Research in Psychology, Hungary. The study was conducted in accordance with the Declaration of Helsinki. Participants received payment (3300 HUF, ca. 10 Euros) or course credit for taking part in the experiment.

### Stimuli

Four pseudowords were used as stimuli: two spoken by a Hungarian native speaker and two spoken by a German native speaker (both were female). The same reiterative pseudoword ‘bébé’ (/be⍰⍰be⍰/) was recorded by each speaker with two different stress patterns: one with stress on the first syllable (/be⍰⍰be⍰/ trochaic stress pattern, henceforth referred to as S1), and one with stress on the second syllable (/be⍰be⍰⍰/ iambic stress pattern, henceforth referred to as S2). Speakers were asked to produce the pseudowords as naturally as they could, and no further software editing was done on the stimuli, except for equalizing the overall loudness (RMS equalization) and applying a 10 ms long onset and offset ramp. The acoustical properties of stimuli are illustrated in Figure 1 and summarized in Table 1.

**Table 1.** Acoustical properties of stimuli.

|  |  | **Hungarian** |  | **German** |  |
| --- | --- | --- | --- | --- | --- |
|  |  | 1<sup>st</sup> syllable | 2<sup>nd</sup> syllable | 1<sup>st</sup> syllable | 2<sup>nd</sup> syllable |
| intensity (dB)<sup>1</sup> | S1 | 82.13 | 66 | 78.79 | 70.8 |
|  | S2 | 75.57 | 80.96 | 74.35 | 78.78 |
| f0 (Hz)<sup>2</sup> | S1 | 225.32 | 202.65 | 213.61 | 215.44 |
|  | S2 | 211.32 | 206.36 | 217.17 | 209.41 |
| duration (ms) | S1 | 360 | 386 | 170 | 403 |
|  | S2 | 350 | 396 | 140 | 433 |

*Note.* <sup>1</sup>: maximum amplitude measured in the whole length of syllable; <sup>2</sup>: maximum f0 measured in the whole length of syllable

**Figure 1.**
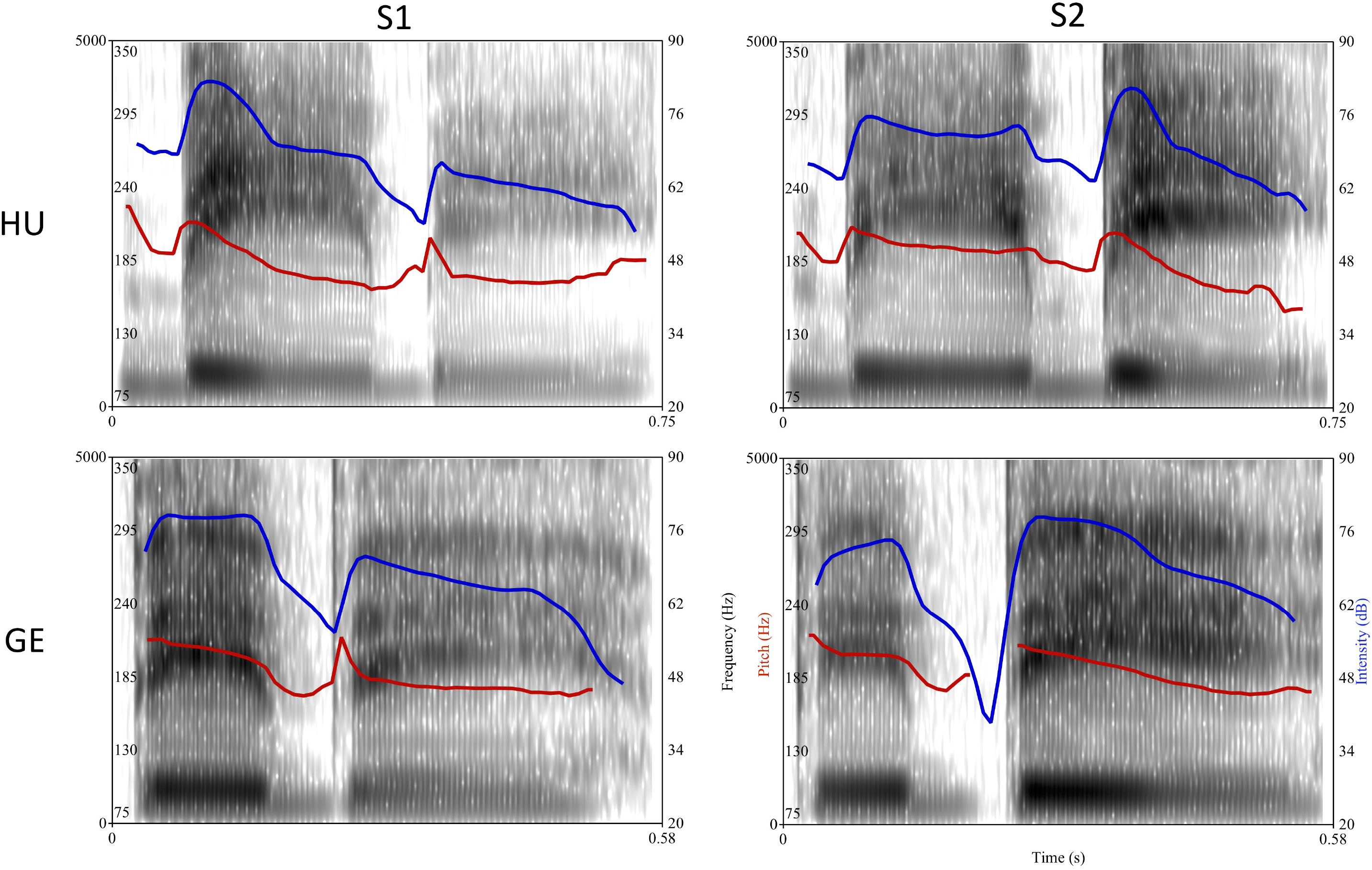
Acoustic characteristics of the stimuli. The figures show the spectrogram, pitch (red line) and intensity (blue line) of the four stimuli. Note the different time axes of Hungarian and German stimuli. *Note*: S1: stimuli with stress on the first syllable; S2: stimuli with stress on the second syllable; HU: Hungarian stimuli; GE: German stimuli.

### Procedure

Stimuli were presented in a *passive oddball paradigm* with a stimulus onset asynchrony (SOA) varying randomly between 850 and 1050 ms. The duration of the S1 stimulus was 746 ms in Hungarian and 573 ms in German, and the duration of the S2 stimulus was 747 ms in Hungarian and 574 ms in German. In our design, we varied language (Hungarian, German), stress position (S1, S2) and role (standard, deviant) yielding *eight different conditions*: the S1 and S2 as a standard stimulus and the S1 and S2 as a deviant stimulus in Hungarian and German, respectively. Stimuli were presented in *four experimental blocks* according to the following pairings: (1) Hungarian S1 standard with Hungarian S2 deviant; (2) Hungarian S2 standard with Hungarian S1 deviant; (2) German S1 standard with German S2 deviant; (3) German S2 standard with German S1 deviant. Participants were presented 666 stimuli in each block with 566 standard stimuli and 100 deviant stimuli (i.e., the probability of the deviant was 15%). Each block began with at least fifteen standard stimuli, and then deviants were separated by at least two standard stimuli. We created a different pseudorandom sequence of stimuli for each block, and within each block, the same sequence of stimuli was used for all participants. The order of the four experimental blocks was counterbalanced across participants. During the EEG experiment, participants were seated in a comfortable chair in an acoustically shielded, dimly lit room. They were instructed to ignore the stimulus sequence presented via headphones (Sennheiser PX 200) while they were watching a silent movie.

After the EEG experiment and the removal of the electrode net, a behavioral discrimination task was administered to check whether the variations in stress position were distinguishable within a language. Namely, a *same-different paradigm* with two blocks was used: one block tested the Hungarian stimuli, another tested the German stimuli. Stimuli were presented pairwise with a SOA of 1000 ms. Forty stimulus pairs were presented in each block: The four possible combinations of stimulus pairs (S1-S1, S2-S2, S1-S2, S2-S1) were tested ten times. The order of blocks was counterbalanced across participants. Participants were required to decide as fast and accurate as possible whether the two stimuli presented as a stimulus pair were the same or different. They had unlimited time to initiate a response for each stimulus pair. The next stimulus pair was presented after the participant gave behavioral response. We used a Cedrus RB-740 response pad (Cedrus Corporation, San Pedro, CA) to record participants’ responses in this task. Assignment of response keys to same/different responses was counterbalanced across participants. The entire experimental procedure lasted about 2 hours, including the application of the electrode net. Both the EEG and the behavioral experiment were written in and controlled by the Presentation software (v. 17.0, Neurobehavioral Systems).

### EEG Recording and Pre-processing

The continuous EEG activity was recorded using the Electrical Geodesics system (GES 400; Electrical Geodesics, Inc., Eugene, OR) and Net Station 4.5.1 software. We used a 128-channel HydroCel Geodesic Sensor Net with saline electrolyte solution (see Figure 2). Electrode Cz was used as a reference and a sampling rate of 500 Hz was applied.

**Figure 2.**
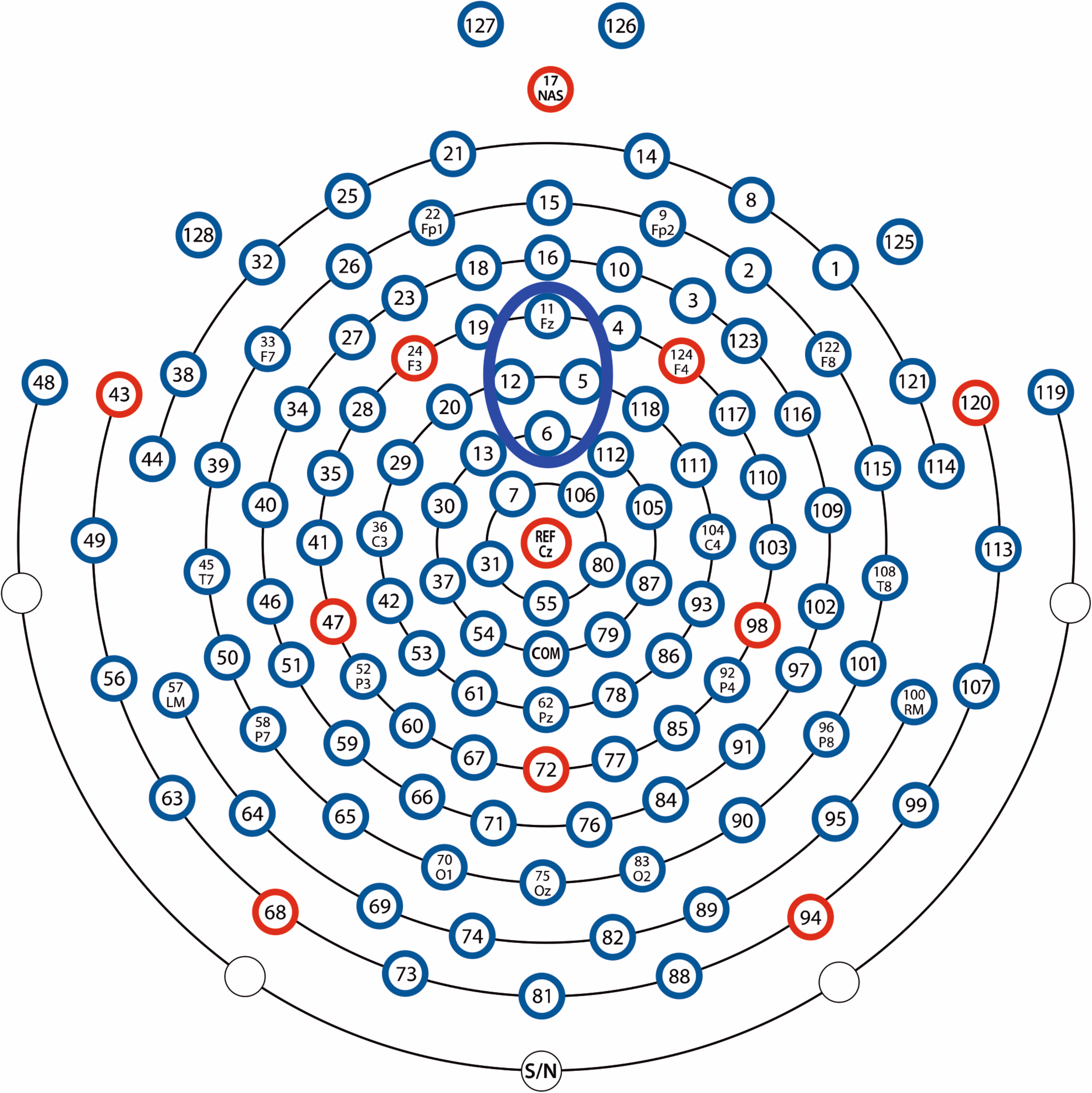
The electrode montage with 128 electrodes used in the experiment. The blue ellipsis shows those electrodes (5, 6, 11, 12) that were pooled together in the fronto-central electrode pool used for statistical analysis.

Data were analyzed using BrainVision Analyzer software (Brain Products GmbH, Munich, Germany) and Matlab 8.5 (MathWorks Inc.). Spline interpolation of bad electrodes was performed if necessary. Zero – three (*M* = 0.61) channels per participant were interpolated. As the first step of pre-processing, EEG data were band-pass filtered offline between 0.3 – 30 Hz (48 dB/oct) and notch filtered at 50 Hz to remove additional electrical noise. Second, we corrected horizontal and vertical eye-movement artifacts and heartbeats with independent component analysis (ICA; Delorme, Sejnowski, & Makeig, 2007). Using ICA, 1-6 ICA components (*M* = 2.93) were removed, then, the channel-based EEG data were recomposed. Third, EEG data were re-referenced to the average activity of all electrodes. After that, the continuous EEG was segmented into epochs. Epochs extended from −100 to 1000 ms relative to the onset of the standard and deviant stimuli, respectively, in each condition. An equal number of epochs (n = 100) were selected for both the standard and deviant stimuli in each condition by analyzing only the standards preceding the deviants. To remove artifacts still present in the data after ICA corrections, we used an automatic artifact rejection algorithm implemented in BrainVision Analyzer software. This algorithm rejected epochs where the activity exceeded +/− 100 μV at any of the electrode sites. Only those participant’s data were included in further analysis where the percentage of rejected epochs was below 30% in any of the conditions. Accordingly, two participants’ data were excluded. The mean number of retained epochs collapsed across all conditions was 95 out of the possible 100 (*SD* = 5.4; range: 72 – 100). After artifact rejection, artifact-free epochs were baseline corrected based on the mean activity from −100 to 0 ms. Finally, the remaining epochs were averaged.

### Data/ERP Analysis

ERP analysis involved a permutation based non-parametric topographic analysis of variance (TANOVA, Murray, Brunet, & Michel, 2008) conducted in Ragu (Koenig, Kottlow, Stein, & Melie-García, 2011) on the individual ERP waveforms. The aim of the analysis was to determine those latency ranges where there are significant differences between the ERP curves related to the effect of interest (see below). The TANOVA allows the quantification of ERP components without selecting time windows arbitrarily (i.e., where the ERP components’ amplitude seems to be maximal or the difference between particular conditions seems to be maximal) or by following previous literature; therefore, it is and unbiased approach (see Brooks, Zoumpoulaki, & Bowman, 2017).

The analysis was conducted separately for the Hungarian and German stimuli. The reason for this was that differences in duration between the stimuli (see Figure 1 and Table 1) made the point-by-point comparison of the ERPs problematic across the languages, as the shorter first syllables in German could potentially elicit ERP components with shorter latencies.

We calculated the TANOVA for topographic dissimilarity (TD) (Koenig & Melie-Garcia, 2009; Wirth et al., 2008), which is a single measure of the distance between two electric field topographies. We assumed that the ERPs to standard and deviant stimuli and to different stress patterns would demonstrate slightly different topographic distributions. The TANOVA was performed with Role (Standard vs. Deviant) and Stress Position (S1 vs. S2) as within-subjects factors. Standard and deviant responses for the same stimuli (i.e., either S1 or S2) taken from different conditions were used in the analysis. This approach enabled us to measure genuine MMN effects as we compared physically identical stimuli within each language differing only in their role (standard vs. deviant) in the experimental blocks (see also Honbolygó & Csépe, 2013; Jacobsen & Schröger, 2003).

We performed 1000 randomization runs and applied a 5% significance threshold on the individual averages during the entire epoch length (no normalization was used). Regarding the main question of this study, we focused on the Stress Position * Role interaction effect, which would show the time range where the difference between deviants and standards is modulated by the stress pattern. As mentioned above, we did not compare effects directly between languages.

## Results

### Behavioral results

One participant’s data in the same-different paradigm are missing because of technical reasons. Thus, behavioral data were analyzed in 29 participants. As Hungarian and German stimuli differed in duration, we did not compare the RT of responses across the different contrasts. Instead, accuracy, defined as the percentage of correct responses, was used as a measure of performance. Responses slower that 2000 ms were excluded from further analysis, thus, accuracy was calculated without these trials. We found ceiling effect on this task as the mean accuracy for the same pairs was 99.48% (*SD* = 1.55%) in the Hungarian block and 98.79% (*SD* = 2.18%) in the German block. For the different pairs, this was 98.45% (*SD* = 3.02%) in the Hungarian block and 99.12% (*SD* = 2.38%) in the German block. Therefore, participants were able to discriminate pseudowords with different stress pattern both in Hungarian and German.

### ERP results

Grand average ERP waveforms to the standard and deviant stimuli and the difference curves split by language and stress position are presented in Figure 3. As it can be seen in the figure, stimuli elicited a complex set of ERP waveforms, starting with positive ERP deflections that went in the negative direction from around 200-300 ms. ERPs to the stimuli in the standard and deviant role started to deviate from each other already from around 100 ms.

**Figure 3.**
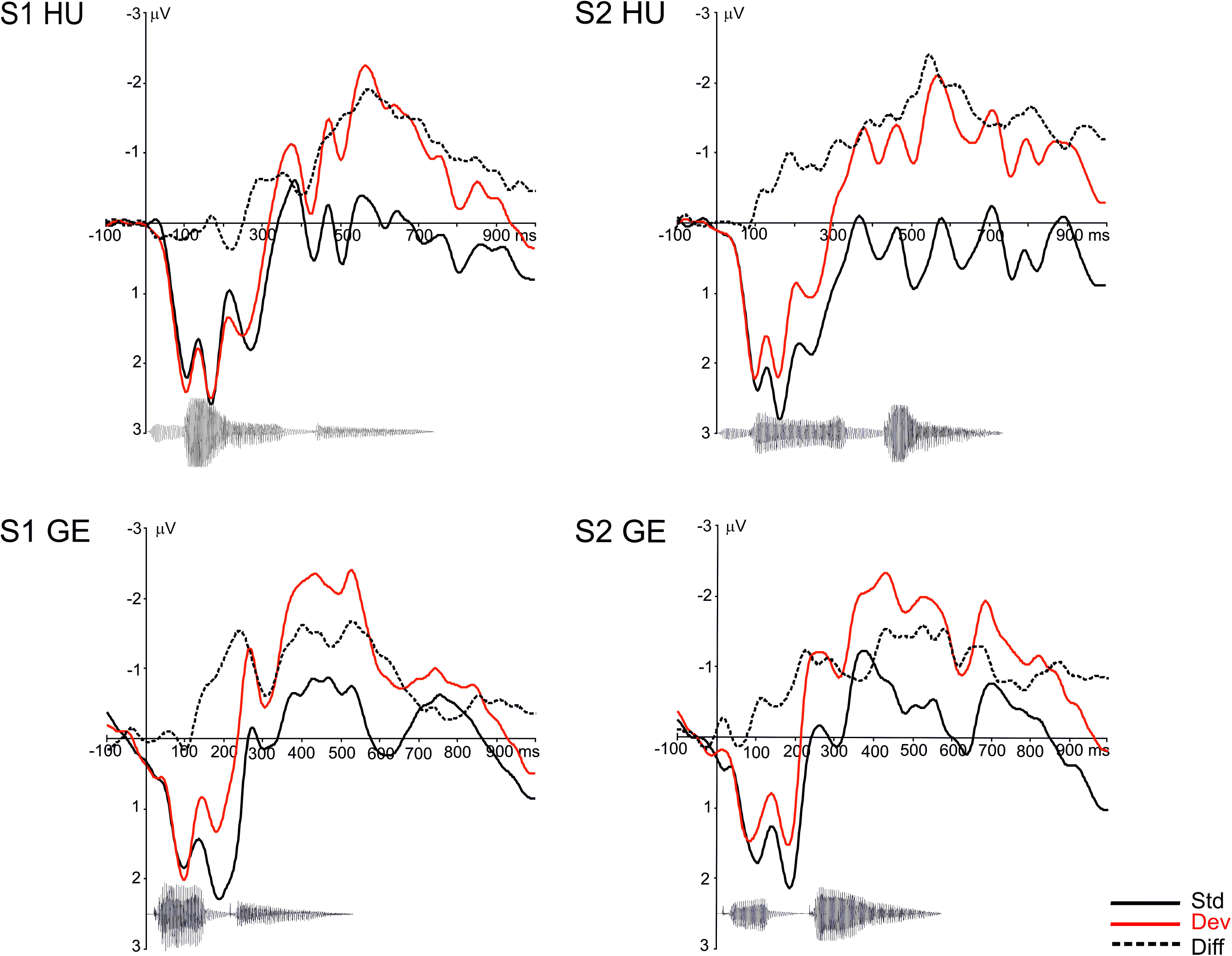
Grand average ERP waveforms split by language and stress position to the standard (Std) and deviant (Dev) stimuli together with the difference wave (Diff) calculated by subtracting the ERPs to standard stimuli from that to the corresponding deviant stimuli. Negativity is plotted upwards. ERPs are plotted for the fronto-central electrode pool. The waveform of the respective stimuli below the ERPs illustrates the temporal relation between the stimuli and ERP responses. *Note*: S1: stimuli with stress on the first syllable; S2: stimuli with stress on the second syllable; HU: Hungarian stimuli; GE: German stimuli.

Difference curves calculated by subtracting the ERPs to the standard from the ERPs to the deviant elicited by the same stimuli are depicted on Figure 4. Difference waves were calculated as follows: (S1 HU) Hungarian S1 deviant minus Hungarian S1 standard; (S2 HU) Hungarian S2 deviant minus Hungarian S2 standard; (S1 GE) German S1 deviant minus German S1 standard; (S2 GE) German S2 deviant minus German S2 standard. The heat maps under the ERPs in Figure 4 show those latency ranges where the TANOVA indicated significant effects (main effect or interaction) for scalp topographies (colors denote the p values). We found a significant Stress position * Role interaction effect only for the Hungarian stimuli between 176-254 ms and 374-418 ms. There was no significant interaction effect for the German stimuli. These two latency windows correspond to two negative ERP deflections, and the significant interaction is due to S1 HU not eliciting or eliciting smaller deflections. As for the German stimuli, these two deflections are also present, but they are not different for the S1 and S2 stimuli. A more detailed presentation of the TANOVA results, and the ANOVA results of the ERP data in the above time windows for both languages separately are presented in the supplementary material.

**Figure 4.**
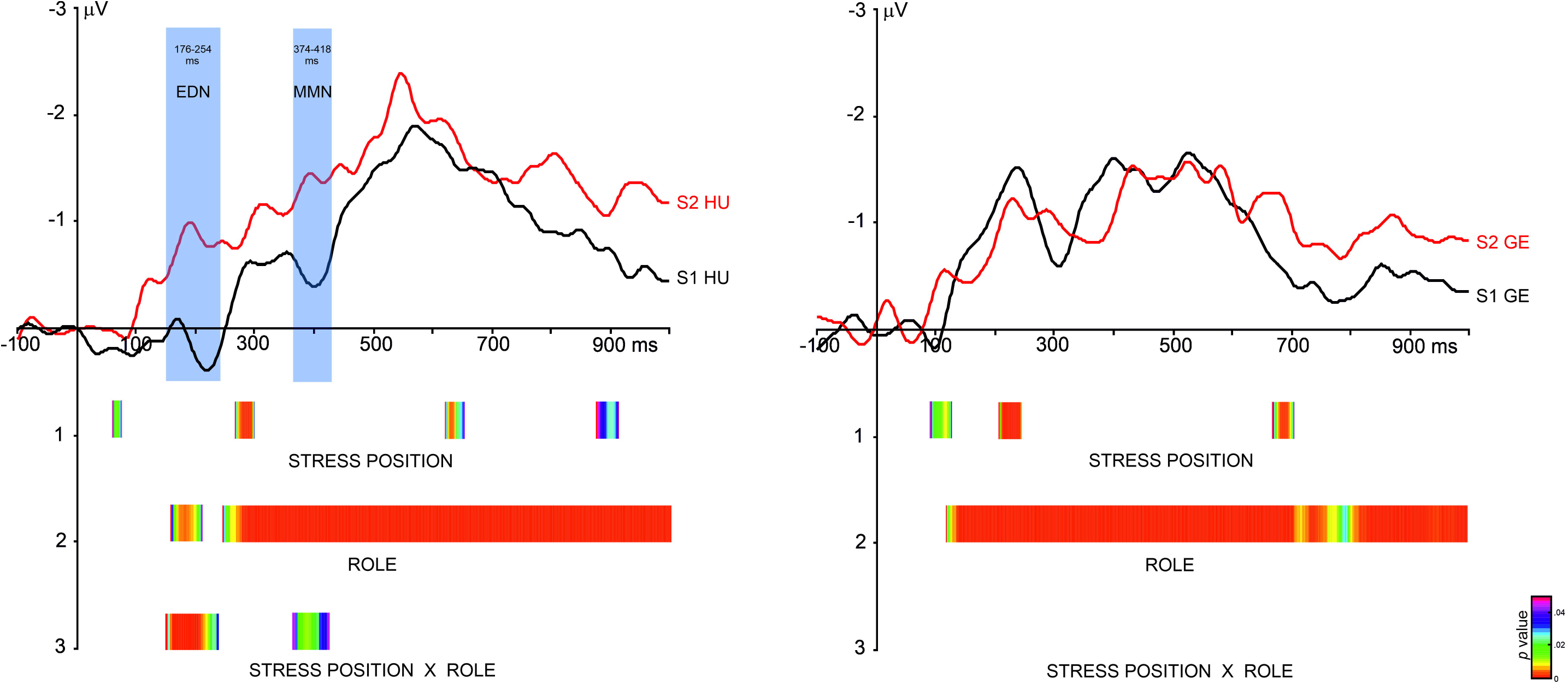
Difference waves in the four different conditions split by language (Hungarian: left side; German: right side) and the TANOVA results. ERPs are plotted for the fronto-central electrode pool. The results of the TANOVA are depicted below the ERPs. The colored intervals show those time windows where the TANOVA indicated significant main effects of Stress Position, Role, and Stress Position * Role interaction. Only those time windows are shown where the effect was significant for at least 10 consecutive time points. The *p*-value of the significance is shown as a colored scale in the range of 0-0.05. Shaded areas show the time windows on the ERPs selected for later analysis (presented in the supplementary material). *Note*: S1: stimuli with stress on the first syllable; S2: stimuli with stress on the second syllable; HU: Hungarian stimuli; GE: German stimuli. EDN: Early Differentiating Negativity.

## Discussion

This study investigated the language-specific nature of word stress representations in a fixed-stress language. We studied the ERP correlates of the processing of disyllabic pseudowords having stress on the first or the second syllable produced by either a native Hungarian or a native German speaker. Using a data-driven analysis approach, we found two time windows that demonstrated stimulus-related effects.

The ERP activity in the first (176-254 ms) and second (374-418 ms) time windows showed a difference between the standard and the deviant stimuli in all conditions except for the Hungarian stress on the first syllable (S1) stimulus. This means that Hungarian listeners detected the difference between the native stress patterns when the stimulus had stress on the second syllable (S2) but not when the stress was on the first syllable. At the same time, they detected the difference between the German stress patterns in both S1 and S2. We interpret these results as evidence for language-specific processing of stress patterns. We suggest that the lack of Hungarian S1 effect means that the legal stress patterns was processed as not being deviant. Based on our earlier results (Garami et al., 2017; Honbolygó & Csépe, 2013; Honbolygó, Kolozsvári, & Csépe, 2017), we assume that the legal stress pattern is processed in relation to long-term stress representations, so called stress-templates. As we argued previously (Honbolygó & Csépe, 2013), the lack of MMN in S1 cannot be explained simply with reference to acoustic differences between the stimuli, as in the case of the trochaic stress pattern, the deviant actually contains more salient acoustic features than the standard.

We suggest that the ERP activity in the first time window is not an MMN component, we rather consider it to be an Early Differentiating Negativity (EDN). The sensitivity of the EDN to feature changes was investigated by Kuuluvainen, Alku, Makkonen, Lipsanen, and Kujala (2016) who studied the discrimination of five syllable features (consonant, vowel, vowel duration, fundamental frequency, and intensity), including both segmental and prosodic phonetic changes in pre-school children. They found that the EDN was elicited around 100-200 ms after stimulus onset by all feature changes except for the vowel duration, and non-speech consonant and f0 changes. Because the EDN was larger for speech than for non-speech stimuli, the authors suggested that this component was related to speech-specific processing mechanisms. They left open the possibility that the EDN could reflect a difference in the neural refractoriness resulting from enhanced N2s for the novel deviant features. In another study investigating stress discrimination (Cunillera, Toro, Sebastián-Gallés, & Rodríguez-Fornells, 2006), the authors found an increased N2 component for the initially stressed stimuli compared with the initially unstressed ones. However, in our study, the EDN component was larger for the initially unstressed syllable compared with the initially stressed one, which suggests that the negative deflection appearing in our study is different from the N2 components. Since data on the EDN component are still scarce, we have to be cautious about the functional interpretation. However, as our data suggest, the EDN is sensitive to both the acoustical information and the regularity of the stimuli.

We considered the ERP deflection in the second time window (374-418 ms) as a genuine MMN component. This component has been previously found in relation to stress processing both in our studies investigating Hungarian language (Garami et al., 2017; Honbolygó & Csépe, 2013; Honbolygó et al., 2017) and in other studies investigating Finnish (Ylinen et al., 2009), German (Weber et al., 2004), and English (Peter et al., 2012) languages. As described above, the MMN component was elicited in the Hungarian S2 and German S1 and S2 conditions but not in the Hungarian S1 condition, indicating that stress-related linguistic information was processed in a language-specific way. The functional difference between the EDN and MMN is currently unclear and requires further investigation. However, based on the temporal relations between the syllables and the ERP deflections (see Figure 3) we can speculate that the EDN and MMN reflect the processing of different information: because its latency is very short, the EDN is probably related to processing the onset of the first syllable, while the MMN may be related to the processing of acoustic-phonetic changes inside the first syllable. Because the second syllable starts around 360 ms, it is unlikely that the MMN would be related to the processing of that.

These results can be interpreted in the AERS model described in the Introduction (Winkler & Schröger, 2015). In our case, the lack of ERP difference in the two time windows for the Hungarian S1 can be explained by assuming that the native stress pattern in the deviant position did not start the updating process because it was fully predicted by the model based on long-term stress-related representations. Thus, in the model, the long-term representations overwrote or overweighed the short-term representations based on the stimulus sequence, in which the deviant stimuli still differed from the standard. In all other cases, the incoming deviant stimuli did start the updating process, because they violated both the short-term (standard vs. deviant) and the long-term (irregular native stress, regular and irregular non-native stress) regularities.

## Summary and conclusion

In summary, we found evidence for the language-specific processing of word stress patterns indicated by the lack of early ERP (EDN and MMN) activities to the native stress pattern in the deviant position. Our results are compatible with a predictive coding model of auditory event representations and suggest that the word stress is processed predictively, based on long-term memory traces activated only by native language input. Further studies are required to extend these findings to languages with variable stress, and to show the limitations imposed by the native memory traces when learning a second language with different stress pattern.

## Acknowledgment

The authors are thankful for Gabriella Baliga, Renáta Szűcs, Lívia Elek for their help in EEG recording and data collection, and Dorottya Gyarmathy and Mária Gósy for their help in recording the stimuli. The study was supported by the Hungarian Scientific Research Fund (OTKA K 119365, PI: V.Cs.), the János Bolyai Research Fellowship of the Hungarian Academy of Sciences (F.H.), and the Postdoctoral Fellowship of the Hungarian Academy of Sciences (to A.K.)

## Footnotes

1 LQ = Laterality Quotient; LQ = −100 means complete left-handedness, LQ = 100 means complete right-handedness.

